# Novel Human Kidney Cell Subsets Identified by Mux-Seq

**DOI:** 10.1101/2020.03.02.973925

**Authors:** Andrew W. Schroeder, Swastika Sur, Priyanka Rashmi, Izabella Damm, Arya Zarinsefat, Matthias Kretzler, Jeff Hodgin, George Hartoularos, Tara Sigdel, Jimmie Chun Ye, Minnie M. Sarwal, for the Kidney Precision Medicine Project

## Abstract

**Background:** The kidney is a highly complex organ that performs multiple functions necessary to maintain systemic homeostasis, with complex interplay from different kidney sub-structures and the coordinated response of diverse cell types, few known and likely many others, as yet undiscovered. Traditional global sequencing techniques are limited in their ability to identify unique and functionally diverse cell types in complex tissues.

**Methods:** Herein we characterize over 45,000 cells from 10 normal human kidneys using unbiased single-cell RNA sequencing. We also apply, for the first time, an approach of multiplexing kidney samples (Mux-Seq), pooled from different individuals, to save input sample amount and cost. We applied the computational tool Demuxlet to assess differential expression across multiple individuals by pooling human kidney cells for scRNA sequencing, utilizing individual genetic variability to determine the identity of each cell.

**Results:** Multiplexed droplet single-cell RNA sequencing results were highly correlated with the singleplexed sample run data. One hundred distinct cell cluster populations in total were identified across the major cell types of the kidney, with varied functional states. Proximal tubular and collecting duct cells were the most heterogeneous, displaying multiple clusters with unique ontologies. Novel proximal tubular cell subsets were identified with regenerative potential. Trajectory analysis demonstrated evolution of cell states between intercalated and principal cells in the collecting duct.

**Conclusions:** Healthy kidney tissue has been successfully analyzed to detect all known renal cell types, inclusive of resident and infiltrating immune cells in the kidney. Mux-Seq is a unique method that allows for rapid and cost-effective single cell, in depth, transcriptional analysis of human kidney tissue.

**Significance Statement:** Use of renal biopsies for single cell transcriptomics is limited by small tissue availability and batch effects. In this study, we have successfully employed the use of Mux-Seq for the first time in kidney. Mux-Seq allows the use of single cell technology at a much more cost-effective manner by pooling samples from multiple individuals for a single sequencing run. This is even more relevant in the case of patient biopsies where the input of tissue is significantly limited. We show that the data from overlapping tissue samples are highly correlated between Mux-Seq and traditional Singleplexed RNA seq. Furthermore, the results from Mux-Seq of 4 pooled samples are highly correlated with singleplexed data from 10 singleplex samples despite the inherent variability among individuals.

## Introduction

Whole-body homeostasis is maintained in large part via filtration, reabsorption, secretion and excretion processes in kidney.^1^ Understanding kidney cell function and its cellular and molecular structure is of fundamental importance in order to understand how to preserve renal health and better predict, diagnose and treat renal disease. Knowledge regarding the transcriptional landscape in kidney has come largely from microarray and bulk RNA-seq, technologies that reflect the average gene expression across thousands of kidney cells, without considering the fluctuations in cell specific gene expression. The recent advent of single-cell RNA-sequencing (scRNA-seq) provides new insights into evaluating kidney sub-structure biological and cellular heterogeneity. Single-cell genomic technologies are now being harnessed for understanding the cellular diversity of different organ systems as part of the Human Cell Atlas initiative^23^, with the deliverable of accurately interpreting cell-specific gene expression data, identify known and new cell types or subtypes involved in disease progression, and follow expression mapped cellular transition states. Single cell studies of the human^4,5,6^ and mouse^7^ kidney have begun to characterize the complexity of renal tissue. Park et al. identified 18 previously known renal cell-types, distinguished sub-types in each known cell-types, and three new cell-types in the mouse kidney^7^. Further efforts are required to fully understand the complexity of the normal human kidney, which has been thought to be composed of more than 25 different cell types, expressing specific proteins, to perform unique functions. Nevertheless, recent scRNA-seq studies on human kidney have often failed to identify known cell types and sub-types, often due to the vulnerability of kidney cells during sample preparation or dissociation method, or variations in batch runs^8,9^, resulting in lack of identification of rare or more sensitive cells. The Chang Zuckerberg Initiative has recently funded a collaborative effort to develop the normal human kidney cell atlas (https://chanzuckerberg.com/science/programs-resources/humancellatlas/seednetworks/a-comprehensive-single-cell-atlas-of-the-human-kidney/) but data from this is not expected until 2020.

To address a rate-limiting challenge for processing kidney tissue for scRNA-seq, we have optimized the methodology for kidney cell dissociation with high cell yield and viability. We present a data analysis pipeline that utilizes unsupervised computational methods to identify 37 unique kidney cell populations, forming almost 100 unique kidney subclusters, from the transcriptomes of over 45,000 human kidney cells. Furthermore, to successfully apply scRNA-seq to study small amount of input kidney tissue from 16-18-gauge needle biopsies, and minimize batch bias and variations, we also present feasibility data on the first ever-successful utilization of a multiplex approach for droplet scRNA-seq (Mux-Seq). The approach of Mux-Seq allows us to transcriptionally profile pooled kidney cells/samples, which can then be computationally deconvoluted to map to kidney cells to individual patients, while significantly reducing the cost, time, and batch effects via pooling cells ^10^. The deconvolution is done by a software tool, Demuxlet, which uses the natural genetic variation among individuals to assign the identity of each cell to the individual source. We show that Mux-Seq has high correlation with the gene expression output obtained by conventional scRNA-seq methods. Therefore, our results provide pilot feasibility of potentially applying scRNA-seq technology on multiple human biopsies at once via Mux-Seq and Demuxlet. Mux-Seq allows the successful identification of different kidney cell clusters in health and disease in a rapid and cost-effective manner. These approaches will allow for an improved understanding of the cellular states and molecular dynamics of kidney health and elucidate the divergence of existing cells and evolution of new cell-subsets in kidney regeneration after injury and disease.

## Methods

### Normal kidney samples

A total of ten human kidney samples were obtained for this study - four from the University of California San Francisco (UCSF) and six from the University of Michigan. All samples were dissected from tumor-free regions of nephrectomy. Samples from University of Michigan were harvested from consented patients by the Tissue Procurement Service as a part of the Kidney Precision Medicine Project (KPMP) consortium (https://kpmp.org/) and were approved as exempted by the University of Michigan Institutional Review Board because they were anonymized. All samples at UCSF were collected under a protocol approved by the University of California San Francisco Institutional Review Board. Informed consent was obtained for useae of samples and data. The samples at UCSF were collected as 18 gauge needle biopsies to mimic closed patient biopsy material. Of the ten samples, there were four males, four females, The gender for two samples were unknown, however, we can infer that these samples were likely male due to their SNP data and the lack of *XIST* gene expression.These 10 samples were all sequenced on single lanes for singleplexed scRNA-seq and 4 samples were also processed separately by the Mux-Seq for comparative analysis with singleplexed runs. The ages of the patients ranged from 57-71 (mean 64.5, SD 5.1). All the tissue samples were processed and stored following the same method. The 4 samples collected from UCSF were processed within one month of tissue collection, whereas the 6 tissues from the University of Michigan were processed at UCSF, when they were received at UCSF ~6 months after local collection. Sample metadata is stored in *<u>Supplementary table S1</u>*.

### Isolation of Cells from Kidney tissue and Single-Cell Preparation

After initial optimizations, we set the following protocol for sample preparation for scRNA-seq. Kidney biopsy tissue of ~7.5 mg (needle biopsy) or nephrectomy dissected tissue (chopped tissue) (was frozen in CryoStor cell cryopreservation media (Cat#C2874, Sigma, St. Louis, MO) and stored in liquid nitrogen until further processing. For processing, tissue was thawed at 37°C and placed in RPMI 1640 medium (Cat#R8758, Sigma, St. Louis, MO) at room temperature for 10min. To isolated cells, the tissue was dissociated by mincing with a sterile razor blade and digesting in RPMI 1640 containing Liberase TL (Cat#5401020001, Sigma, St. Louis, MO) and DNase I (Cat#4536282001, Sigma, St. Louis, MO). After 30 min, the cell suspension was filtered through a 70um cell strainer into cold RPMI medium supplemented with 10% of fetal bovine serum (FBS). The filtrate containing cells, was centrifuged at 300g for 10 min at 4°C, the cell pellet washed and resuspended with ice-cold PBS with 0.4% BSA. Cell-viability was determined using a mixture of ethidium bromide and orange acridine. For Mux-Seq each pool contained 5000 resuspended live-cells from each of four individual patients. Cells were resuspended in PBS + 0.04% BSA to a final cell concentration of 1000 cells/μL for library preparation.

### DNA Isolation and Exome Sequencing or SNP array analysis

Genomic DNA was isolated from kidney biopsies using QIAamp DNA Mini Kit (Cat# 51104, Qiagen, Hilden, Germany), obtaining ~30ng of DNA from 5000 cells. DNA was sent to the UCSF Institute of Human Genetics Core for genotyping using SNP arrays (OmniExpress Exome kit, Illumina).

### Single-cell RNA-seq Library Preparation and Sequencing

Libraries for single-cell RNA-seq were prepared using the 10X Single Cell Immune Profiling Solution Kit according to the standard manufacturer protocol. Briefly, single cell suspension, 10X barcoded gel beads and oil were loaded into Chromium Single Cell A Chip to capture single cells in oil droplets (Gel Bead-In-Emulsions, GEMs) at a targeted cell recovery of 4000-8000 cells. Full length cDNA libraries were prepared by incubating GEMs in a thermocycler. Following reverse transcription and cell barcoding in droplets, emulsions were broken and cDNA purified using Dynabeads MyOne SILANE followed by PCR amplification (98°C for 3 min; 12-16 cycles of 98°C for 15 sec, 67°C for 20 sec, 72°C for 1 min; 72°C for 1 min) for 3’ gene expression sequencing library construction. Amplified cDNA was fragmented and end-repaired, double sided size selected with SPRIselect beads, PCR amplified with sample indexing primers and repeat double-sided size selected with SPRIselect beads. Single-cell RNA libraries were sequenced on the Illumina NovaSeq S2 to a minimum sequencing depth of 50,000 reads/cell using the read lengths 26bp Read1, 8bp i7 Index, 91bp Read2.

### Single cell and multiplexed single cell RNA sequencing: generation of data matrix, quality control, and preprocessing

Data was processed via the 10X Chromium 3’ v2 platform. Data matrices and barcode information were generated using the 10X Cell ranger version 2.1.1 software, aligned to the GRCh38 transcriptome. After data generation, barcode-matrix preprocessing was performed to remove cells of low quality for downstream *in silico* analysis. We applied two-phases of cut-offs for barcodes kept in the analysis. Initially, we examined the distribution of cells across clusters with mitochondrial reads per cell <50% and >500 genes per cell. Next, upon determining that these thresholds did not affect our clustering, we expanded the final threshold to keep cells with <80% mitochondrial content and >200 genes per cell in an effort to maximize our cellular population. Additionally, cells with >6,000 genes per cell were excluded to eliminate the likelihood of bias introduced by the presence of doublets. The final data matrix for the singleplexed scRNA-seq dataset included 33,694 genes across 45,411 cells.

### In Silico Demultiplexing

For validation and correlation of the Mux-Seq technique with traditional singleplexed scRNA-seq, 4 kidney samples (also used in the singleplexed analysis) were used. Cells from each of the four kidneys were pooled and run in duplicate wells on the same 10X platform. Applying the same QC metrics as above, the resulting data matrix contained 33,694 genes across 7,574 cells. The Demuxlet tool was then applied to these cells to determinate the sample origin as well as doublet probabilities using at least 50 unique SNPs per sample.

### Data Analysis

The first phase of analysis was performed in R using the Seurat^11^ version 2.3.4 package. Gene expression data was first normalized by multiplying each expression by 10,000 then log-transforming. Then the top 2,000 most highly variable genes were used for downstream dimension reduction. The data was scaled by regressing out the effects of the number of unique molecular identifiers (nUMI) per cell as well as percentage mitochondrial transcripts (%mito). Linear dimension reduction was done by computing the top 50 principal components. These 50 principal components were then used for non-linear dimension reduction, shared nearest neighbor graph construction and Louvain clustering, and visualization via tSNE. The resolution parameter of the Seurat tSNE function, which controls the number of clusters produced, was set to 1.0. For the 2^nd^ level subclustering analysis, after cell barcodes were annotated to cell type, the data matrices for each cell type were individually put through the pipeline again to develop the subclusters. Clusters were numbered in the decreasing order of cluster size as determined by the number of cells in the cluster.

In order to determine the identity of individual cell populations, differential expression analysis was done. Average expression of each gene was calculated for each cell population (cluster), then logged fold change was determined by taking the average expression of a single gene in one cluster against all other clusters. FDR-corrected p-values were calculated using the Wilcoxon rank-sum test. Genes with a minimum log fold change value of 0.5 and expressed in at least 25% of the cluster were included. Then, genes were assigned a marker score as calculated by the fold change of expression multiplied by the ratio of percent prevalence of the gene in cluster versus all other clusters. Once a cluster’s identity was determined using a list of well-known marker genes (Supplementary Table S2), this process was repeated with each broad population of cell types for subclustering to further explore heterogeneity within cell types. For functional profiling of cell population clusters, gene set enrichment analysis (GSEA) was performed using the R bioconductor package clusterProfiler version 3.10.1.

The Monocle 2^12^ tool version 2.10.1 package in R was used for pseudotime computation and cell trajectory analysis of a cell type. Genes were selected for ordering and significance testing for association with pseudotime if they were expressed in at least 10 cells and had a mean expression value of at least 0.05. The data associated with this project is available at the Data Lake of the Kidney Precision Medicine Project (www.kpmp.org).

## Results

### Single cell profiling and unbiased clustering of human kidney cells reveals 37 distinct kidney cell clusters

Using kidney tissue collected from nephrectomies by wedge resection (n=6) and renal biopsy (n=4), we optimized a protocol for single cell isolation and established a pipeline for droplet scRNA-seq (*Figure 1*). Next, we successfully utilized a technique of multiplexed single cell sequencing (Mux-Seq)^10^ to pool cells from 4 different patients and compare the average gene expression of all genes to that obtained from parallel sequencing of individual patient samples by singleplexed scRNA sequencing. Mux-Seq allows the pooling of cells from multiple individuals in one microfluidic run and leverages the genotypes of each sample to assign it to each individual cell using a bioinformatic demultiplexing tool, called Demuxlet, which is part of the Mux-Seq workflow.

**Figure 1.**
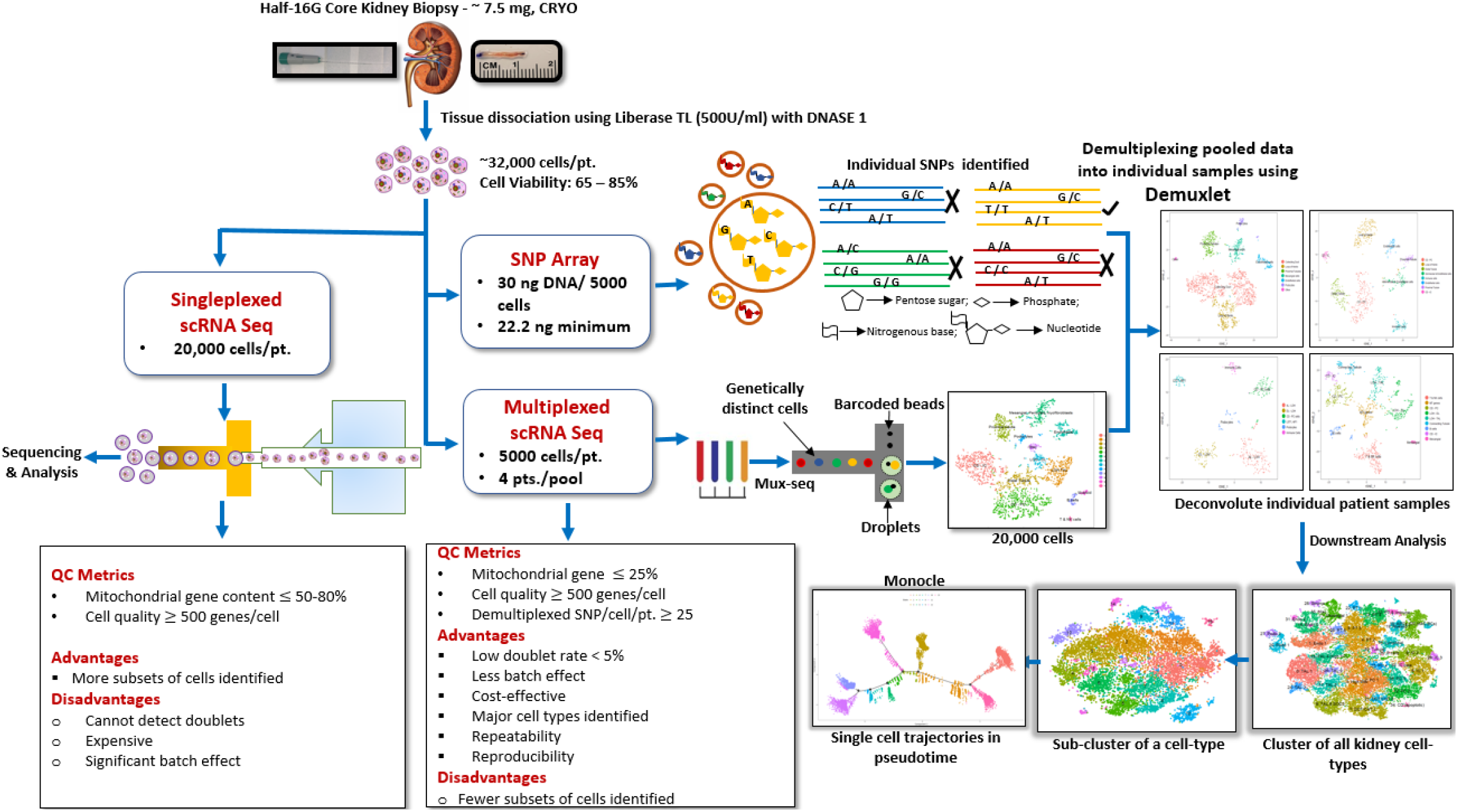
Overview of kidney tissue processing for droplet based scRNA-seq. A pipeline indicating optimized protocol for cell isolation and data processing are shown for singleplexed as well as multiplex RNA-seq.

Demuxlet was able to leverage SNP array data to identify the sample origin of the 7,574 pooled cells, as well as identify doublets using a minimum of 50 SNPs unique to each sample, resulting in an overall detected doublet rate of 6.2% (*Figure 2A*). As with the singleplexed method, all major kidney cell types were identified by Mux-Seq (*Figure 2B*) and direct comparison of average expression of all genes between the scRNA-seq versus Mux-Seq showed high concordance (Spearman’s rho = 0.97; *Figure 2C*) as well has high concordance of expression between selected canonical kidney cell marker genes (*Figure 2D*). Together, these results show a high reproducibility of cluster mapping with our Mux-Seq protocol despite the smaller input of tissue and cells.

**Figure 2.**
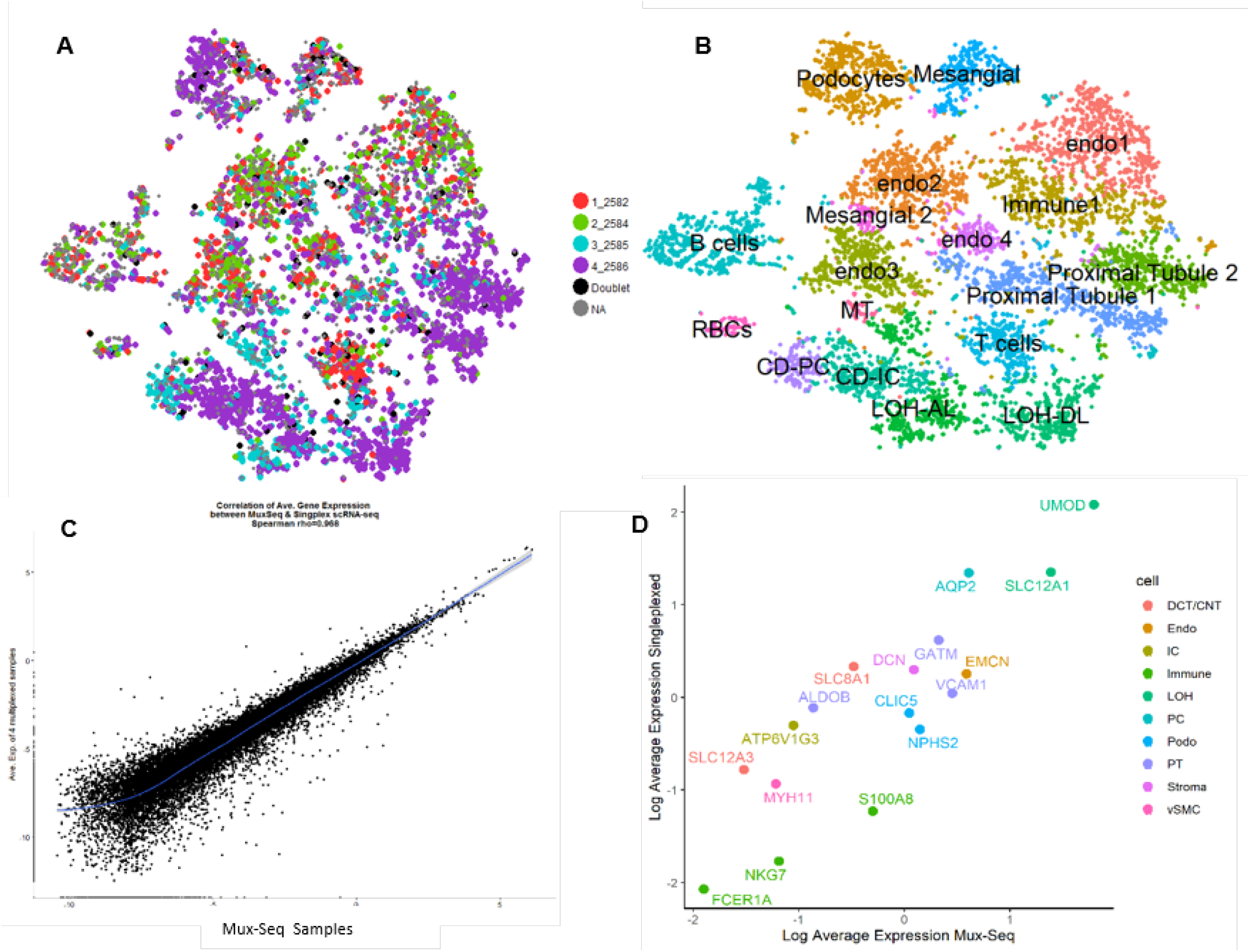
Demuxlet data and comparison to Singleplexed data. (A) Shows a tSNE plot of 7,574 cells from nephrectomies from 4 separate individuals from the overall singleplexed dataset pooled together for sequencing (Mux-Seq). Points are colored by individual, detected doublet, and unmatched cell to sample origin (NA). (B) Shows the tSNE from (A) colored by all major kidney cell types. The scatterplot in (C) shows high correlation between the Singleplexed and Mux-Seq methods by average gene expression of each cell. The scatterplot in (D) also shows high correlation (Pearson r= 0.94) between the average expression of selected top marker genes were all kidney cell types for both the Singleplexed and Mux-Seq data. Cluster labels are abbreviated as follows: Glomerular endothelial cells (Endo), Loop of Henle ascending and descending limb (LOH – AL, LOH – DL), Principal and Intercalated cells of the Collecting Duct (CD – PC, CD – IC), Red Blood Cells (RBCs), cells unidentifiable due to only gene markers being mitochondrial reads (MT).

The pipeline was established for both one sample per microfluidic run (singleplexed scRNA-seq) and pooled samples in one microfluidic run (Mux-Seq). 37 distinct kidney cell clusters (ranging from 50 to 3,196 cells/ cluster) totaling 45,411 single cell transcriptomes were identified from the 10 kidney samples (*Figure 3*). All but one cluster contained cells from multiple patients (*Supplementary Figure S1*). Differential gene expression analysis was used to generate cluster-specific marker genes. Based on the unique gene expression profile of each cluster (*<u>Supplementary Table S3</u>*), we identified cell types representing each renal compartment as well as resident and infiltrating immune cells. In this way, endothelial, stromal/interstitial, podocytes, mesangial, proximal tubule, thick ascending limb of the loop of Henle, distal tubule, connecting tubule, collecting duct, and immune cells were identified and annotated (*Figure 3B*) with the top gene markers for each cluster displayed in figure 3C. Three clusters (3, 30 and 35) were unidentifiable to any distinct known parent cell category, based on differential gene expression analysis of known cell-specific genes. Interestingly, for several cell types we identified more than one cell populations: 3 endothelial, 2 podocyte, 6 proximal tubule, 9 collecting duct and 4 immune cell clusters.

**Figure 3.**
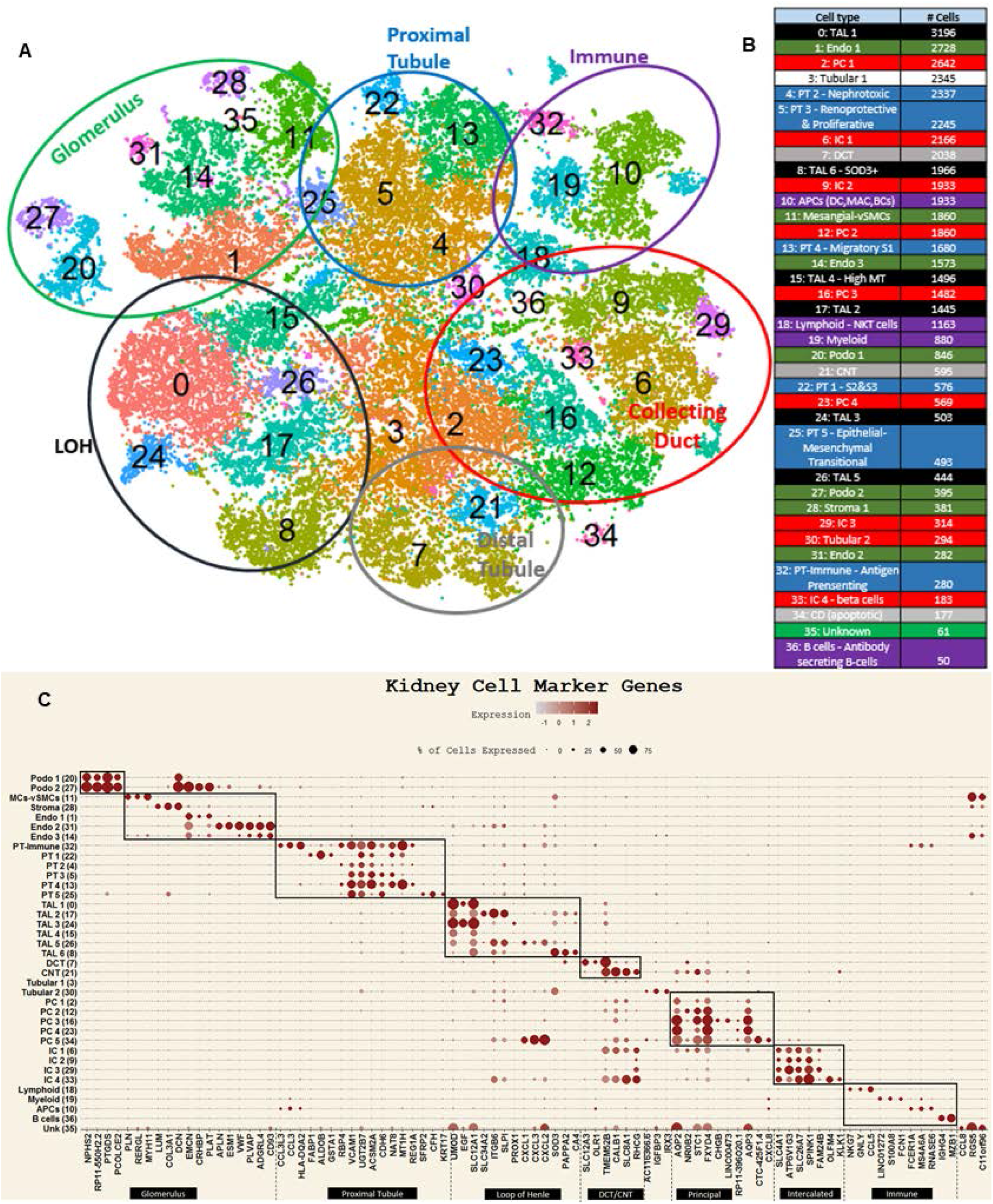
Single cell transcriptomic analysis of human kidney reveals 37 distinct cell populations. (A) 45,411 single cell transcriptomes from 10 nephrectomy tissues were analyzed. Unsupervised clustering resulted in 37 distinct cell populations shown in a tSNE plot. (B) Table listing clusters shown in (A) with cell type annotations and number of cells in each cluster. Each cluster is color coded to reflect the cell type in (A). (C) Dot plot of average gene expression values (log scale) of top 3 unique differentially expressed markers and percentage of cells expressing these markers in each cluster.

Next, we performed several quality control analyses to validate our cluster map. Most of our clusters contain cells from multiple individuals of both genders resulting in low batch effects (*Supplementary Figures S1A, B*). Furthermore, we correlated our expression results with single nuclear RNA sequencing data from human kidney^4^ and found good correlation in the average expression of the 731 most highly variable genes common to both datasets (*Figure 4*) further strengthening the robustness of our data. Finally, we explored the association of transcriptomic mitochondrial read content with kidney function by gene set enrichment analysis (*Figure 5*).

**Figure 4.**
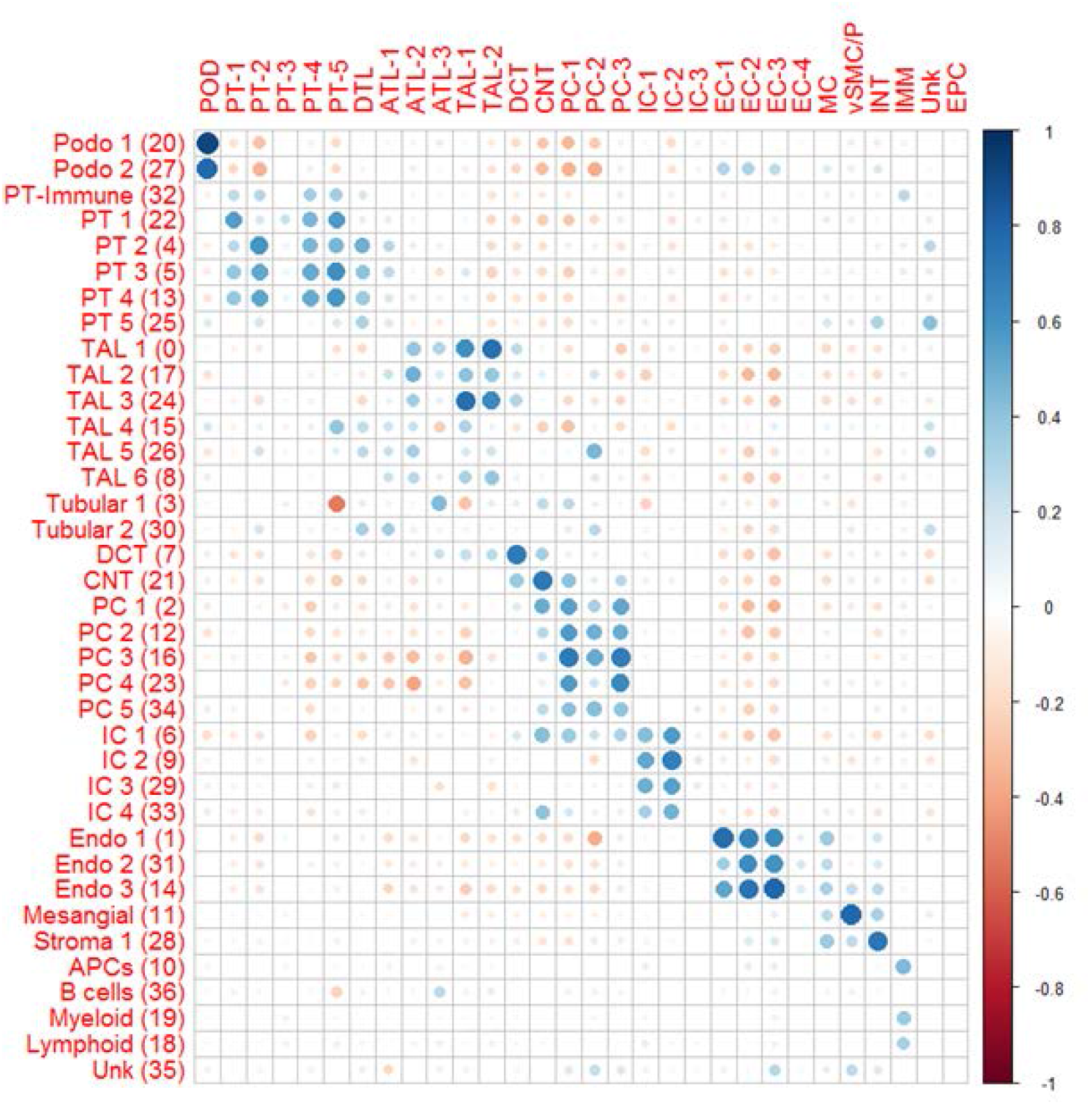
Heatmap showing high correlation between scRNA-seq and snRNA-seq data. Correlation of average gene expression of common variable genes with cell clusters of snRNA-seq data of the kidney from Lake et al.^4^ with our results. Larger, darker blue circles denote higher Pearson correlation. Asterisks are placed next to clusters of low correlation (i.e., r<0.40).

**Figure 5.**
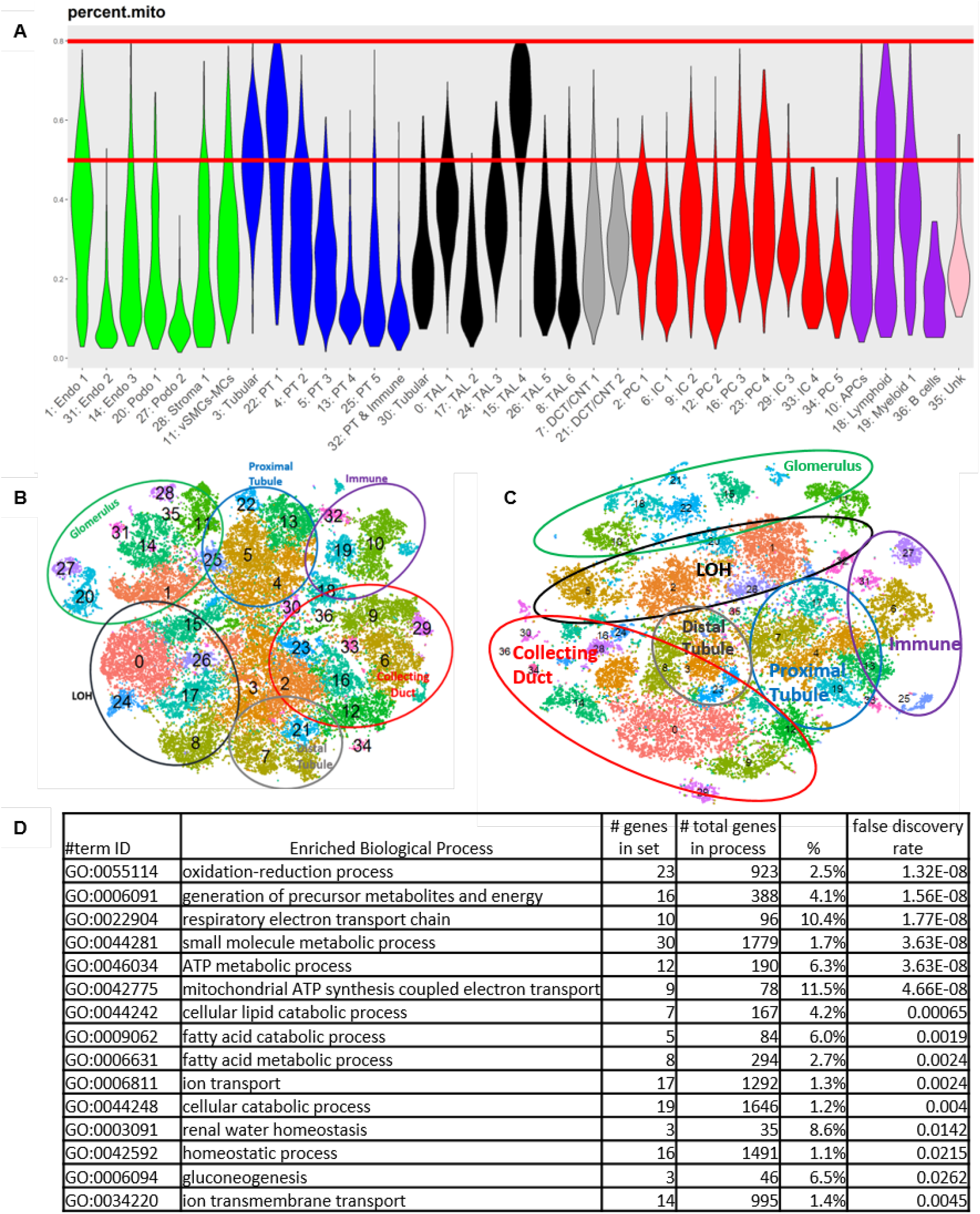
Mitochondrial gene content correlates with kidney function, not stress responses. (A) Shows violin plots of mitochondrial transcriptome content by cluster. Two red lines are present to compare cells from clusters lost due an 80% versus 50% maximum threshold. Only clusters 3 (tubular), 15 (TAL), and 21 (PT) had a mean %mito content greater than 40%. The mean %mito at the 80% cut-off is 31% while the mean %mito at the 50% cut-off is 26%. (B) and (C) are tSNE plots of clusters identified with the mitochondrial thresholds of 80% and 50%, respectively. (D) Displays results of gene ontology testing for significantly enriched biological processes. These were derived from the 125 genes that were positively correlated with mitochondrial gene percentage based on Pearson’s correlation (FDR adjusted p-value < 0.05).

### Characterization of Glomerular cells reveals podocytes as the main site of injury in proteinuric diseases with underlying genetic mutations

Cluster analysis identified all major glomerular components including podocytes, glomerular endothelial cells (Cluster 1/Endo 1), mesangial cells, and vascular smooth muscle cells (Cluster 11/Mesangial-vSMCs). Besides expressing the canonical markers of glomerular endothelial cells, Endo 1 also expressed *IGFBP5* (Insulin-like growth factor binding protein) at high levels (*Figures 2A, 2C*). Using well-known podocyte markers, NPHS1 and NPHS2, we identified two podocyte clusters: cluster 20 (Podo 1) and cluster 27 (Podo 2) (*Figures 2A, 2B*). Podo 1 is the bigger cluster and displays high expression of genes with a functional role in glomerular filtration. This cluster is also enriched for genes involved in podocyte actin cytoskeletal organization and regulation-hallmark of podocyte filtration function. Hence, we identified this cluster as the native podocyte population. The Podo 2 cluster has fewer cells and expresses markers of podocyte cellular regulation and responses to chemical stimuli. However, Podo 2 cluster was predominantly derived from a single patient possibly reflecting changes associated with any underlying treatments the patient might have received. We named this population as responder podocytes.

We further investigated the podocyte cluster assignments by scRNA-seq with the expression of 29 known genes associated with monogenic inheritance of proteinuria in humans across cell types. We found that we were able to detect the expression of 25 out of 29 genes in our dataset. While most genes (21 out of the 25 genes) were mainly expressed in podocytes confirming that proteinuria is mainly a defect of podocytes, we also detected a high expression of certain genes in other cell types (*Supplementary Figure S2A*). *ACTN4* (α-actinin 4) showed significant expression in almost all clusters but was highest (>3 fold) in podocytes (*Supplementary Figure S2A*). To ensure that the expression of *ACTN4* is not an artifact of cellular dissociation and processing, we validated ACTN4 protein expression using the Human Protein Atlas^13^. Immunohistochemical staining of ACTN4 is consistent with our data showing expression in both glomerular and tubular cells (*Supplementary Figure S2B*). Park et al. had previously shown that podocyte dysfunction is the principal reason for proteinuria based on the mouse homolog expression of the genes associated with nephrotic syndrome^7^. However, in humans there seems to be a contribution of other cell types as well as demonstrated by their high expression in non-podocyte cells.

### Differentiated Proximal Tubule cells retain their ability to proliferate after injury

Unbiased clustering of all the transcriptomes resulted in 6 distinct populations of proximal tubule cells (cluster 22/PT 1, cluster 4/PT 2, cluster 5/PT 3, cluster 13/PT 4, cluster 25/PT 5 and cluster 32/PT 6) (*Figure 2A and 5A*). These clusters did not separate into S1, S2 and S3 segments based on marker genes only. This is consistent with previous studies suggesting a lack of discrete transition from one segment to another^8^. Cells in PT 1/ cluster 22 are enriched in metabolic and detoxification activity predominantly carried out by S2-S3 segment (*Figure 6C*). Interestingly, PT 1 is the only cluster that expresses a lactate transporter (*SLC5A12*) and phosphoenolpyruvate carboxykinase (*PCK1*) enzyme-the main control point for regulation of gluconeogenesis (*<u>Table S3</u>*). As lactate is the principal precursor for gluconeogenesis in humans^14^, our results suggest that PT 1 cells are largely responsible for renal glucose production. PT 2 cells are enriched in genes associated with response to heavy metals and negatively regulate cell growth (*Figure 6D*).PT 3 cells are enriched in genes associated with stimulation of transcription and translation of coding as well as non-coding genes (*Figure 6E*). PT 4 has high expression of metallothioneins, which might be a protective mechanism in the kidney with ageing (*Figure 6F*). PT 5 consists of proximal tubule epithelial cells, which are undergoing mesenchymal transition such as becoming fibroblasts/myofibroblasts. They are associated with increased extracellular matrix formation/organization, and activation of adhesion-mediated signaling pathways (*Figure 6G*). Epithelial cells in PT 6 plays a role in inflammatory and immunoregulatory processes (*Figure 6H*). However, this cluster is almost entirely derived from cells of single patient and might reflect individual peculiarity. Interestingly, all PT clusters but PT-1 express low but significant levels of *CD24*, *CD133*, and *VIM* (vimentin) - markers of putative epithelial stem cells in the human kidney^15^. It has been suggested that differentiated epithelia in humans undergoes spontaneous injury followed by homeostatic repair. This repair is achieved by reversible dedifferentiation and proliferation of fully differentiated tubular cells^15^. In order to test the variation of cellular differentiation states within PT cells, we used the unsupervised algorithm Monocle^12^ that captures single cell variation to order cells in “pseudotime” by progress through differentiation. This allows us to recognize multiple cell fates stemming from a single progenitor cell type. Our trajectory analysis revealed a continuum of cells branching at one point into two different fates arising from a root in state 3 that consists of a mixed population of cells from all clusters (PT 2, 3, 4, 5) expressing markers of putative stem cells(*Figure 7A, B, and C*) At the branch point these cells spilt into two branches: state 2 comprised of cells mainly from PT 3 cells that still retain the putative stem cell like properties and state 1 mainly composed of PT 1 cells that are involved in the canonical tubular activities with some contribution from PT 2 and PT 3.

**Figure 6.**
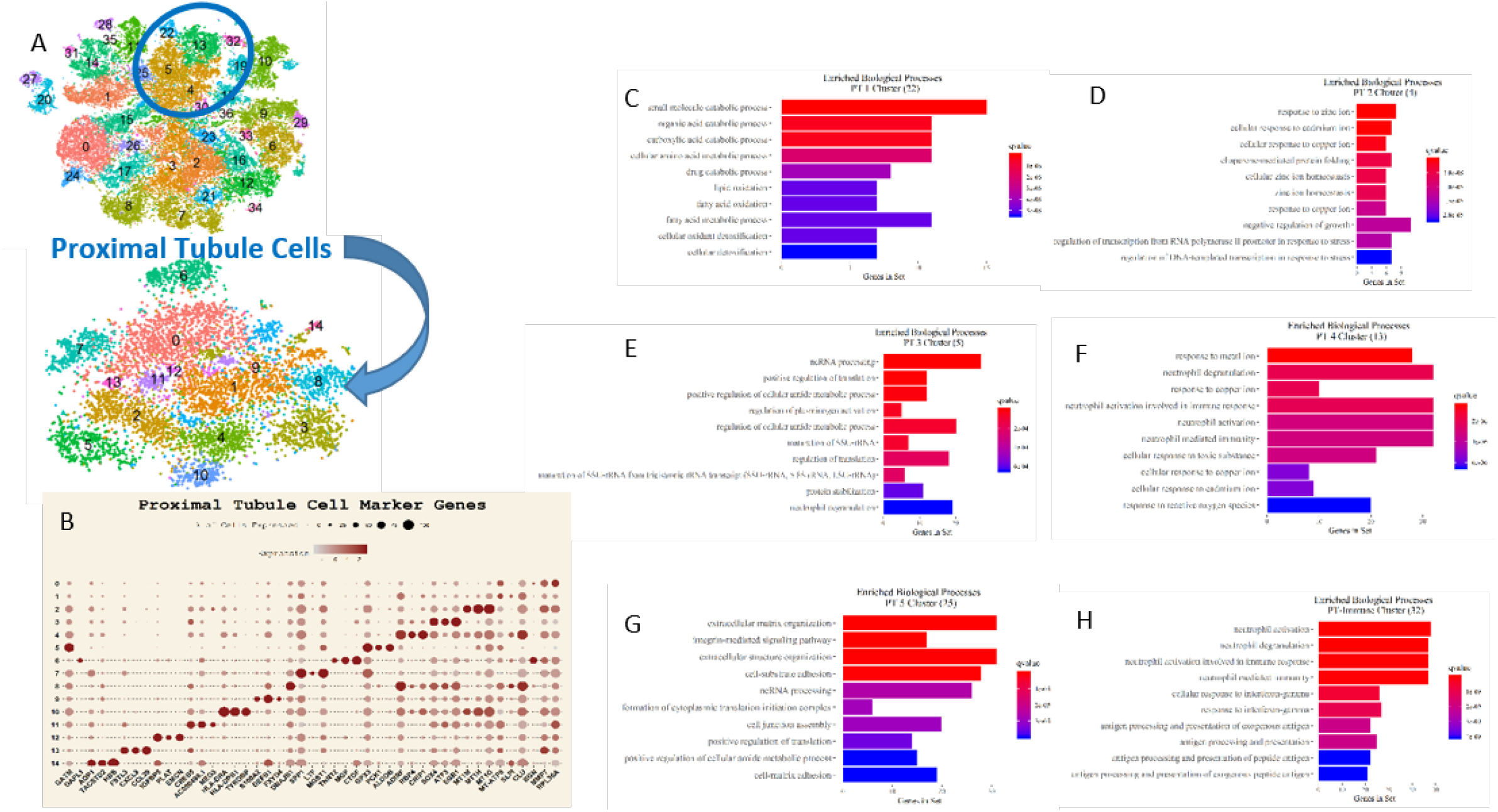
Characterization of proximal tubule cells. (A) Represents the in silico isolation of the 6 parent cluster of PT cells that are then put through the clustering pipeline again resulting in 15 PT subclusters. (B) The top 3 markers for each sub cluster shown in (A) by fold change. (CH) Bar plots show gene set enrichment analysis of each of the 6 PT clusters. The top 10 unique enriched biological processes (BP) were chosen to show functional contrast. The x-axis shows how many genes from the PT cluster’s differentially expressed gene (DEG) list are involved in the BP.

**Figure 7.**
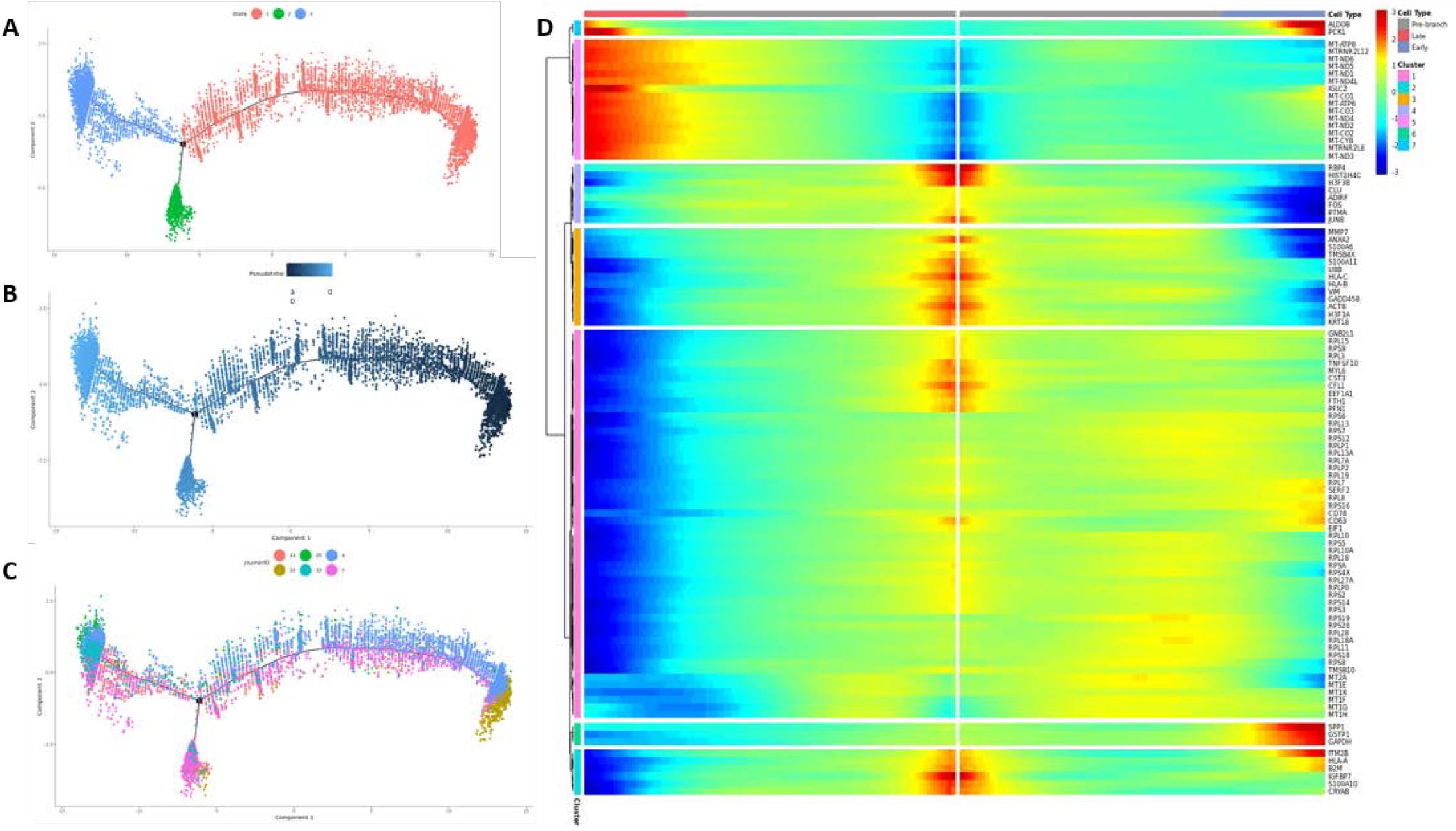
PT cell trajectory analysis using Monocle pseudotime. (A-C) The cell trajectory plots show all PT cells ordered by pseudotime and colored by state (A), pseudotime (B), and mapped to original parent cluster (C). (D) Heatmap of the 100 genes most significantly associated with pseudotime. Pseudotime is displayed in the x-axis, with darker red indicating a later pseudotime state and blue indicating an earlier pseudotime state. Genes on the y-axis were grouped into 7 clusters in a supervised manner based on similar patterns of pseudotime trajectory.

Gene expression analysis of early (fully differentiated quiescent PT cells), pre-branch (dedifferentiated and proliferating PT cells) and late stage (regenerated and quiescent PT cells) shows that expression of mitochondrial genes (*Figure 7D* - heatmap cluster 5) is significantly reduced in pre-branch stages of dedifferentiation and displays highest expression in late-stage mature PT cells. Ribosomal gene expression is significantly higher in early and pre branch pseudotime states compared to that of later stages (heatmap cluster 1) suggesting cell growth, proliferation, and differentiation. As expected, genes associated with stress and injury were significantly higher in pre branch stages (heatmap clusters 2, 3, and 4) reflecting spontaneous injury and dedifferentiation while gluconeogenic genes (*Figure 7D* - heatmap cluster 7) were highly expressed in early and late stage PT cells. These findings strongly support the earlier reports suggesting that human proximal tubule epithelia in healthy state undergo spontaneous injury followed by homeostatic repair. Furthermore, the proliferative capacity of renal proximal tubule after injury involves bulk of differentiated epithelial cells.

### Characterization of Loop of Henle Cells

Cells of the Loop of Henle were the second most abundant cell type (*Figure 2A*). As mostly the cortex was sampled, cells were overwhelmingly from the Thick Ascending Limb (TAL). There were six TAL clusters totaling 9,050 cells, with the largest cluster overall being a TAL cluster. Each of these clusters were labeled because of their significant differential expression of the well-known TAL genes *UMOD* (uromodulin) and *SLC12A1* (NKCC2) (*Figure 2C*). However, our second largest TAL population of nearly 2,000 cells (cluster 8) was unique in that they were *SLC12A1^+^/UMOD^-^*. The top 3 markers of this cluster were *SOD3, PAPPA2,* and *CA4* (*Figure 2C*). Superoxide Dismutase 3/Extracellular Superoxide Dismutase (SOD3/EC-SOD), the most specific marker to this cluster, has been shown to be reno-protective and is characterized by a heparin-binding domain that can anchor the protein to the endothelium and the extracellular matrix^16^. Pappalysin 2 (PAPPA2) has been previously reported in the TAL^17^..

### Characterization of collecting duct cells reveals plasticity of principal cells

Differential gene expression analysis identified 9 populations of cells as collecting duct based on marker genes (*Figure 2A, B*). Four clusters (clusters 2/PC 1, 12/PC 2, 16/PC 3, and 23/PC 4) are enriched in genes consistent with their physiology as principal cells however PC 1, PC 2 and PC 3 also express intercalated cell marker ATP6V1 at lower but significant levels. Intercalated cells separated into four clusters (clusters 6/IC 1, 9/IC 2 and 29/ IC 3)., IC 1, IC 2 and IC 3 appear to be Type A Intercalated cells from OMCD due to the co-expression of *SLC26A7* and *SLC4A1* and pathway analysis showed enrichment of genes involved in the collecting duct acid secretion. We were not able to identify a distinct type B but cluster 33/IC 4 expressed markers of type A as well as type B intercalated cells (cluster 33/IC 4). Cluster 34 is mainly derived from one patient and shows markers of apoptosis. This cluster may reflect the subpopulation of cells in the individual patient affected by the underlying disease or treatment. The plasticity of collecting duct subpopulations was evident when we analyzed their cellular states using monocle. We found that even in adult kidney majority of principal cells retain the ability to differentiate into intercalated cells (*Supplementary Figure S3*). To investigate the transition of principal cells into intercalated cells at single cell resolution, we performed a cell fate trajectory and pseudotime analysis using the aforementioned Monocle tool. Our trajectory analysis revealed a continuum of principal cells with four distinct branch points showing a root corresponding predominantly to the principal cell populations 1, 2 and 3 (state 2, 4, 5, 6, 7, 8 and 9). A branch point arose from these clusters forming state 10 and 11. State 10 is a mixture of IC 1 and PC 1/2 while state 11 is PC cells only. Interestingly, IC 1 is the only intercalated cell cluster that showed low but significant levels of principal cell marker AQP2 expression. The two other branches (state 3 and 12) are composed of intercalated cell clusters IC1/2 and 1 respectively. Another branching point at the end stage led to states 1 and 13 composed mainly of intercalated cells and principal cells respectively. While further sampling is necessary, our results suggest that PC and IC cells undergo cellular transitions in adult human kidney similar to that has been previously described in mouse.

### Characterization of Immune cells

Using known markers, we identified 4 immune cell clusters in our dataset. Cluster 10 is enriched in markers specific to antigen presenting cells (APCs), including markers of dendritic cells, macrophages, and B Cells thus making it a pool of different APCs. Cells of cluster 18 were of lymphoid origin. This cluster is composed of cells from almost all the patients and expresses markers for both, NK cells and T-cells. Cells of myeloid origin were present in cluster 19, which is also enriched in *LST1*-marker for monocyte differentiation^18^ (*Figure 2A, 2C*). We also identified an activated B-cell cluster (cluster 36).

### Subclusters of renal cells identify novel cell populations

Further comparative analysis of cell populations representing renal segments resulted in identification of smaller subsets of cells expressing unique genes suggesting that not all cells of the same type are in the same dynamic and temporal state. Interestingly, we were able to find at least one subcluster in each cell type that has high expression of inflammatory genes suggesting that there are injured/stressed cells present in healthy tissue as well (*<u>Table S4</u>*).

We demonstrate the workflow to further investigate these subclusters in Figure 6. We separated the proximal tubule cells into 15 subclusters (*Figure 6A*) and characterized based on top genes (*Figure 6B*). A proliferative PT cell population (enriched in ribosomal proteins, and markers for cell growth, proliferation, and development^19,20^) was identified in subcluster 0. PTs with expression for protease inhibitors in subcluster 1 maybe more relevant in kidney disease, as these cells are found to be elevated in minimal change disease and HE4, a gene in this subcluster, is known to be a potential biomarker for CKD diagnosis^21–23^. Subcluster 2 is a subset of PTs with enriched in metallothionein-encoding subunit genes including *MT1G, MT1H, and MT1M*; known to be specific to the S1-segment, genes that can be impacted by increasing renal age^24,25^. Moreover, we found this family of genes to be significantly upregulated in the patient with unilateral ureteral obstruction (*Supplementary Figure S4*). Subcluster 3 has cells that are proliferating or developing and undergoing cell spreading, with upregulation of *EGR1* and *SOX4* in subcluster 3, both are required for the normal progress through mitosis and renal development^26,27,28,29^. Upregulation of LIM/double zinc finger protein family members was observed in subcluster 4, which plays a role in cell repair and motility^30,31^. Subcluster 5 has increased expression of genes that play an important role in the metabolic process including expression of Aldolase B, Fructose-Bisphosphate (*ALDOB*) and may belong to the more metabolically active S2-S3 renal segment. Subcluster 6 expresses markers for the proximal tubule epithelial cells, which are undergoing mesenchymal transition including, *CTGF*, *TAGLN*, and *LGALS1*^32,33^. Among others, this subcluster has increased expression of two unique genes, TNNT2 and DAPL1, which have never been reported to be expressed in the kidney before, and appear to be novel PT markers, expressed in approximately 50% and 43% of PT-cells, as opposed to only 2% and 1% of non-PT cells, respectively. Subcluster 7 is enriched in genes playing a role in in renoprotection, based on their role in detoxification from oxidative stress and glucose reabsorption^34^ (*LTF, MGST1, and MAP17*). Subcluster 8 has PT cells with a stress response phenotype, with increased expression of several Heat Shock Protein Family members (*DNAJB1, HSPA6, and HSPA1B*). Subcluster 9 is enriched in PT cells with active transport, with high levels of Na -K-ATPase, water and calcium-binding channels including, *FXYD4, S100A2, AQP2, and CALB1*. Subcluster 10, is enriched in PT-cells playing a role in inflammatory and immunoregulatory processes, with strong signal from HLA genes, IFNγ and neutrophil transmigration^35^. Subcluster 11 has a higher expression of non-protein coding genes and ribosomal pseudogenes, known to be involved in kidney apoptosis^36^. Subcluster 12 consist of PT-cells that has upregulated genes that negatively regulate cell adhesion to the extracellular matrix due to *EMCN* expression^37^. Subcluster 13 is enriched in genes associated with chemokine-mediated signaling pathway; 83% of these cells expressed *CCL20* as opposed to 4% of non-PT cells. This subcluster may also play a role in PT related immune surveillance as *CCL20* has been shown to play a nephroprotective role during AKI, both by decreasing tissue injury and facilitating repair^38^.

Similarly, podocytes separated into six subclusters expressing the traditional podocyte markers with additional cell specific markers (*Supplementary Figure S5A, B*). We found one subcluster (subcluster 3) that showed enrichment of genes involved in sodium ion transport and maintenance of transmembrane electrochemical gradient (Table S3). Approximately 44% of cells in this cluster expressed *SLC12A1* (NKCC2), a well-established marker for the TAL (*Supplementary Figure S5C*). To ensure that the expression of these atypical genes is not an artifact of cellular dissociation and processing, we tested *SLC12A1* (NKCC2) expression in renal biopsy by immunohistochemistry (IHC). IHC confirmed low and restricted but significant expression of NKCC2 in podocytes (*Supplementary Figure S5D*). These results were also supported by the Mux-Seq data (*Supplementary Figures S5E and S5F*).

The largest heterogeneity of different cell subsets was seen in cells expressing known collecting duct genes. A total of 17 subclusters were seen and differences in the different subclusters relate to the pivotal functions of the collecting duct relating to solute and water transport. Subcluster 1, 2, and 9 expressing PC cell markers *AQP2, FXYD4* and *STC1*. Subclusters 4 and 5 are alpha intercalated cells expressing markers *ATP6V0D2* and *SLC26A7*^4^. There was clear evidence of collecting duct cells that formed the Loop of Henle (LOH); *UMOD* is a known LOH marker. And is co-expressed in subcluster 13 with *SLC12A1*, another known LOH marker. Other collecting duct markers show co-expression of highly-regulated genes such as *TMEM* and *FAM24B*, which have previously thought not to be expressed in the kidney^13^. Additionally, like in other renal cell sub-structures, some of the collecting duct cells (subclusters 11, 12 and 14) also show high expression of immune genes such as TNF, interferon, and cytokine signaling pathways.

Subclustering of immune cells provided higher resolution and allowed us to identify distinct components of resident as well as infiltrating populations of immune cells in kidney. We identified 14 immune cell subclusters and analyzed them based on their marker genes (*<u>Table S5</u>*) and pathway analysis to identify novel markers of immune cells. These clusters show highly varied immune cells, in different stages of development and differentiation, such as early and late stage B cells, NK, dendritic cells, and developmental stages of monocytes and macrophages. Subcluster 1 was enriched in dendritic cells and monocyte markers^39^. APCs with MHC class II were represented in subcluster 3 which also expressed the dendritic cell marker *CST3*^40^. Subcluster 4 expressed *CD69*, a marker for tissue resident T cells^41^ and *IL7R* which is present on NK cells^42^. Subclusters 5 and 6 are the only cells that express *LST1* (MHC class III) suggesting APC maturation^18^. Subcluster 9 expressed NK cell markers *GZMA*^43^, *NKG7*^44^, along with *CD3, CD8A, CD69*^41^, and *INFG*^45^ suggesting that these are tissue resident CD8+ cytotoxic T cells and NK cells. Subcluster 13 was mainly derived from cluster Subcluster 9 was very interesting as it identified an exclusive tissue resident macrophage population. Subcluster 13 was mainly derived from cluster 36 and was enriched in markers for memory B cells^46^. Subcluster 14 carries an active T-cell marker, *CD96*^47^ along with classical T-cell markers *CD3D* and *IL2RG*.

## Discussion

Though the technology for high throughput profiling of a large number of cells by droplet scRNA-seq has substantially improved, the applicability of scRNAseq technology towards creating a renal (and other complex tissue) cell atlas in health and disease have been hampered by significant limitations. These limitations include: availability of clinical patient tissue, high costs of running multiple samples by scRNASeq, and run/batch variations -- all of which can be readily addressed by applying the Mux-Seq methodology. In a proof of principle approach, we present the efficacy of the first Mux-Seq data on human kidney samples, and propose it as an ideal solution for handling limited human samples without loss of single-cell tissue biology. We were able to identify >90% of doublets which is consistent with the predicted doublet identification rates with as few as 50 SNPs per cell.

In this study, we provide protocols that optimize tissue processing for scRNA-seq from limited amounts of human kidney tissue to mimic sample volumes that would normally be available during interrogation of clinical tissue samples from patients. We report high yield and viability after renal tissue dissociation to robustly classify single-cell heterogeneity of 10 kidney samples, provide a comparative analysis of robust kidney single-cell biology from Mux-Seq and singleplex scRNA-seq and highlight the Demuxlet methodology approach for data deconvoluton by Mux-Seq. We report a finding of higher mitochondrial content in the human kidney and provide rationale for evaluating single-cell data at different mitochondrial thresholds to interrogate the depth of heterogeneity of the human kidney at the single-cell level. We provide a discussion of inclusion of multiple human kidney cell populations, more than previously classified, with an aim to harness a comprehensive transcriptomic map of the kidney. We use unsupervised, unbiased computational methods to resolve 37 unique cell populations from the transcriptomes of over 45,000 human kidney cells and describe the potential of 100 single cell sub-clusters in the kidney, expanding on our previous appreciation of kidney cellular heterogeneity.

Our understanding of single-cell biology of the human kidney also highlights species (human versus murine) associated similarities and differences in kidney disease pathogenesis. We show that while the expression of monogenic kidney disease genes is suggested to be restricted to a single cell type in mouse, in human kidney, disease pathogenesis is more complex and for a number of monogenic diseases, more than one kidney cell type appears to be involved (*Supplementary Figure S2*).

Using cell trajectory analysis we also attempt to address cell lineage relationships in the kidney, highlighting the collecting duct and proximal tubule epithelia, and again observing species specific differences in injury repair. The proximal tubule lineage during injury repair^48^ appears to be different in humans versus rodent models of tubular injury, specifically relating to variations in the transcriptional regulation of CD24, CD133 and vimentin positive populations of differentiated proximal tubules in humans at baseline, and in rodents only at injury^49,15^. More than 11,000 human collecting duct cells have been analyzed in this study, and transitional cells are noted in the human collecting duct, consisting mostly of PC cell type that can transition into IC cells. The presence of transitional cells in collecting duct in rodents has been described before^15,50^.

As our study performed single-cell rather than single-nuclear sequencing, it was able to identify most of the immune cell types reported in the kidney^51,52^, and immune-cell subclusters identify tissue resident as well as infiltrating immune populations in kidney. Immune cells do not appear to be artifactually overrepresented in this dataset, which has been suggested as a limitation of scRNA-seq methodology by others^53–55^. Our initial clusters only separated into cells of lymphoid vs myeloid origin, APCs and activated B-cells. Subclustering however, led to identification of markers of tissue residence, such as CD45 (*PTPRC*), *IFNG*, and *CD69*^51^ in our lymphoid subclusters. Tissue resident macrophages have been well defined in rodents but there have been technical limitations to identify them in humans^52^. Recent single cell RNA sequencing efforts have contributed to putative markers that can be used to identify this population, and tissue resident macrophages could be identified in the normal kidney, with *APOE* being a putative novel marker to distinguish this population. Immune cell clusters were also identified that expressed MHC and Th17 positive cells^56^; correlations of these expressing cells with age are suggested but this study lacks power for definitive analysis of any clinical association. Enrollment of larger sample numbers as part of the recent initiative to develop a normal human kidney single cell atlas across different age groups (https://chanzuckerberg.com/science/programs-resources/humancellatlas/), will provide the needed numbers to address age specific variations of kidney single cell variations,

A limitation of this study is that though we find that there are new sub-populations of renal cells (Demuxlet did not assign any of these cells to be doublets), such as new podocyte, proximal tubular, thick ascending limb and collecting duct clusters that express new marker genes, restricted tissue availability prevents us from performing detailed validation of these novel marker genes by immunohistochemistry or *in situ* hybridization, and only limited staining validation data is available in this study (*Supplemental Figure S4*). With advances in spatial imaging and with coordinated tissue access planned through KPMP, these validation studies will be performed subsequently, using advanced techniques such as CODEX^57^ and MIBI^58^. To further improve on doublet detection, we propose to take advantage of recently developed tools such as kBET to assess Mux-Seq’s ability to further reduce batch effects^59^.

We also understand that in a tissue of high metabolic and energy demands such as the kidney, a higher mitochondrial burden is expected, when compared to single-cell datasets from PBMC. We address the issue of single cell dissociation mediated introduction of stress-induced transcriptional artifacts^55^, and discuss that while stress-response and mitochondrial genes are present in our data, they appear ubiquitously and produce one likely artifactual cluster of 61 cells; cluster 35, labeled Unk (Unknown) and only 3 clusters (tubular clusters 3, 15, 22) have a mean mitochondrial content greater than 40% (Figure 5). We optimized the selection of mitochondrial threshold analysis in this human kidney by comparing clustering with different thresholds (see methods) and have regressed out the effects of mitochondrial content, along with other QC metrics (*Supplementary figure S6*). The mean percent mitochondrial transcript content in our data was 31% (standard deviation of 16%) similar to that obtained by Wu et al using the singleplexed scRNA-seq. Several studies have proposed the use of mitochondrial to ribosomal gene content ratio for removing dead/dying cells from analysis^60^. We did not observe any predilection for loss of cell viability in different kidney regions, as there was similar mitochondrial to ribosomal gene content ratio across different cell populations. Given that the kidney is the second most mitochondria-rich organ behind the heart^61–63^, our findings suggest that the kidney cells are highly metabolically active and a higher mitochondrial threshold than used in other tissues^9^ may actually be necessary to capture the entire spectrum of renal cell function. Additionally, as shown in *Figure 5*, genes correlating significantly with cellular mitochondrial content often correlate and relate to genes with roles in normal kidney function, rather than other genes indicative of cell stress.

In summary, we have successfully established a pipeline to interrogate limited volume clinical specimens using the Mux-Seq technology and discovered potentially novel kidney cell subtype markers via droplet single-cell RNA sequencing with an aim to explore the sub-cellular heterogeneity in the normal human kidney. To enable transition of scRNA-seq technology to the patient bedside we need to ensure compatibility with clinical samples where tissue quantities by core needle biopsies have to be shared with pathology and are thus necessarily limited. We have ensured feasibility of getting reliable and biologically relevant single-cell data from complex human samples by using tissue sizes in the needle biopsy range. Mux-Seq allows us to establish an optimized pipeline to process and analyze multiple human kidney biopsy samples simultaneously in a cost effective and time efficient manner. Lowering cost and batch-effects by pooling samples while retaining valuable biological information is an important advancement toward the goal of one day making scRNA-seq standard practice in the diagnostic workup of kidney biopsies.

## Author Contributions

M.S. conceived the study. A.W.S. developed and performed the computational analysis. All authors contributed to cluster analysis. M.S., A.W.S. and S.S. created figures. S.S. and I.D. processed samples and performed experiments. M.K and J. H. provided some of the tissue samples from the University of Michigan. G.H. and J.C.Y. assisted in data processing and analysis. M.S., A.W.S., P.R., S.S., A.Z, and T.S. wrote the manuscript. J.C.Y. and M.S. directed the study.

## Acknowledgements

This work was funded by the Kidney Precision Medicine Project (KPMP) Consortium. Juliane Liberto, Parhom Towfighi, Reuben Sarwal and Andréa Alice da Silva assisted in collection of data on the UCSF tissue samples.

## Supplemental Material

Table S1. Sample metadata.

Table S2. Known kidney cell markers used for cluster annotation.

Table S3. Top gene markers for each kidney cell cluster.

Table S4. Top gene markers for each of the 100 subclusters.

Table S5. Top gene markers for each Mux-Seq cluster from figure 2.

## Supplementary Figures

**Figure S1.**
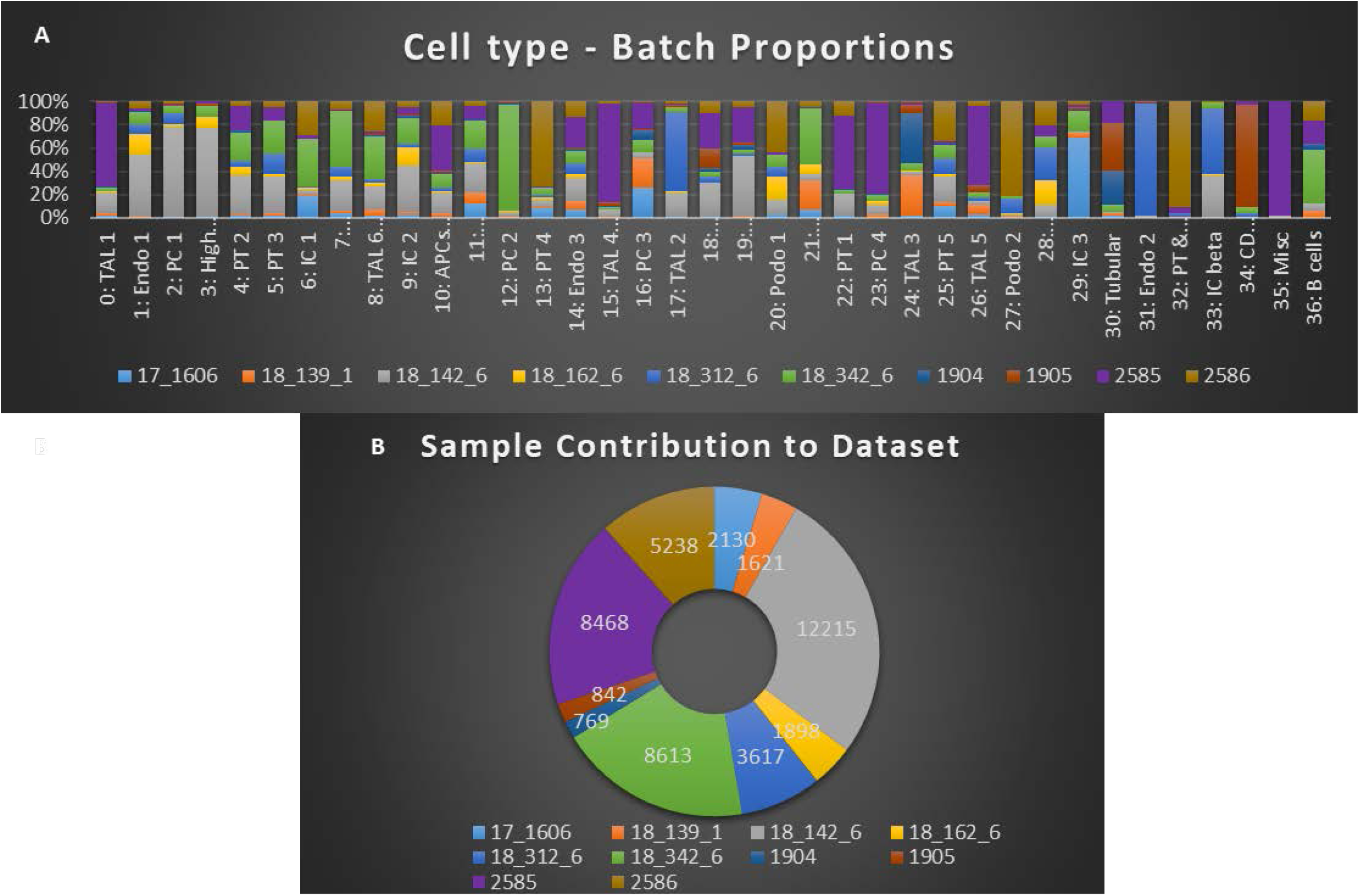
Distribution of cells from each individual shows minimal batch effect. (A) Bar graph shows the distribution of cells from each individual in 37 clusters identified. (B) Proportion of cells from each individual contributing to the entire data set.

**Figure S2.**
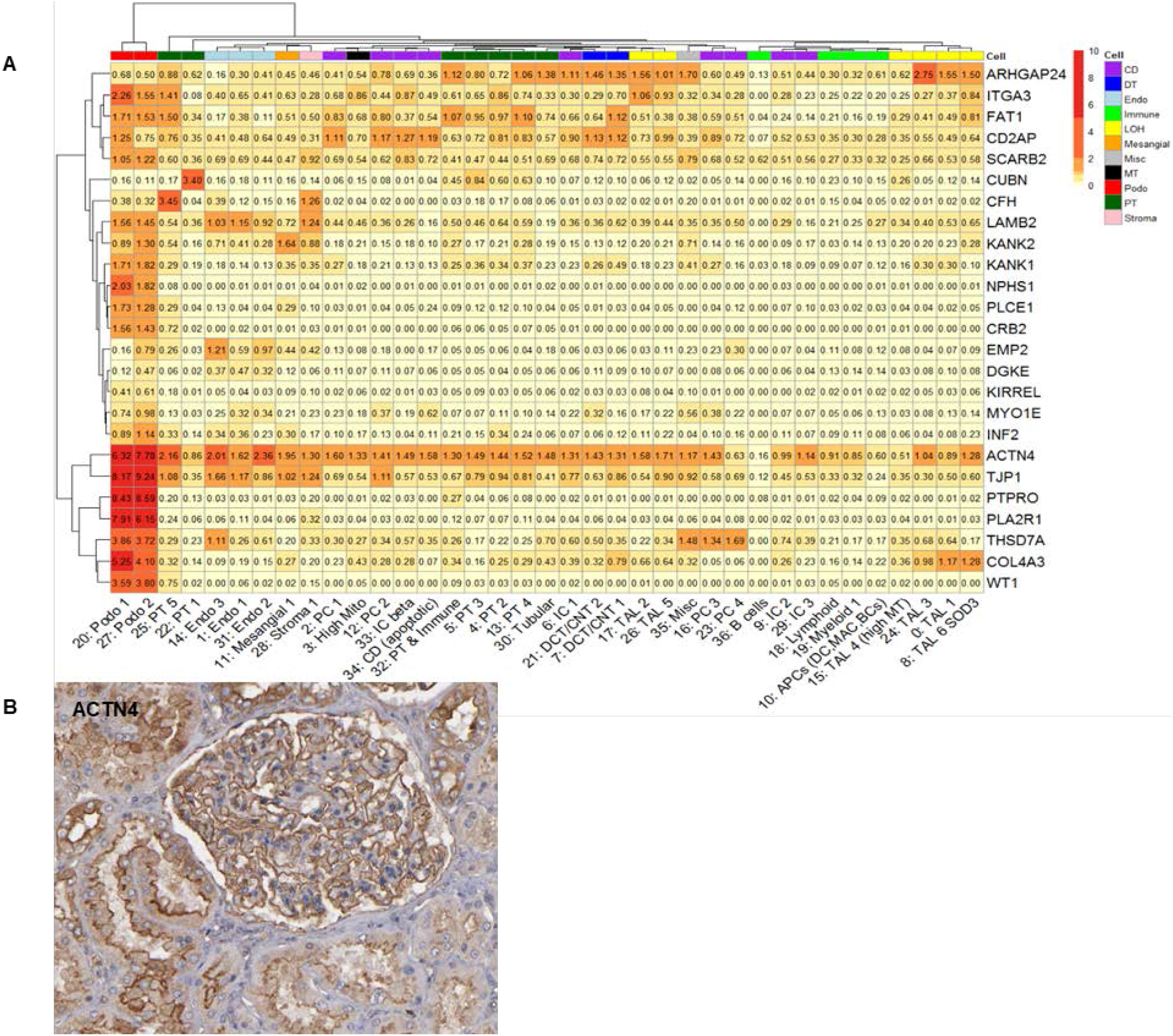
Heatmap showing high expression of genes associated with nephrotic syndrome. (A) Average gene expression of 25 genes associated with inherited forms of nephrotic syndrome were compared across 37 clusters identified and represented in a heatmap. (B) Protein immunostaining (Human Protein Atlas) of *ACTN4* (α-actinin 4) in the glomerulus.

**Figure S3.**
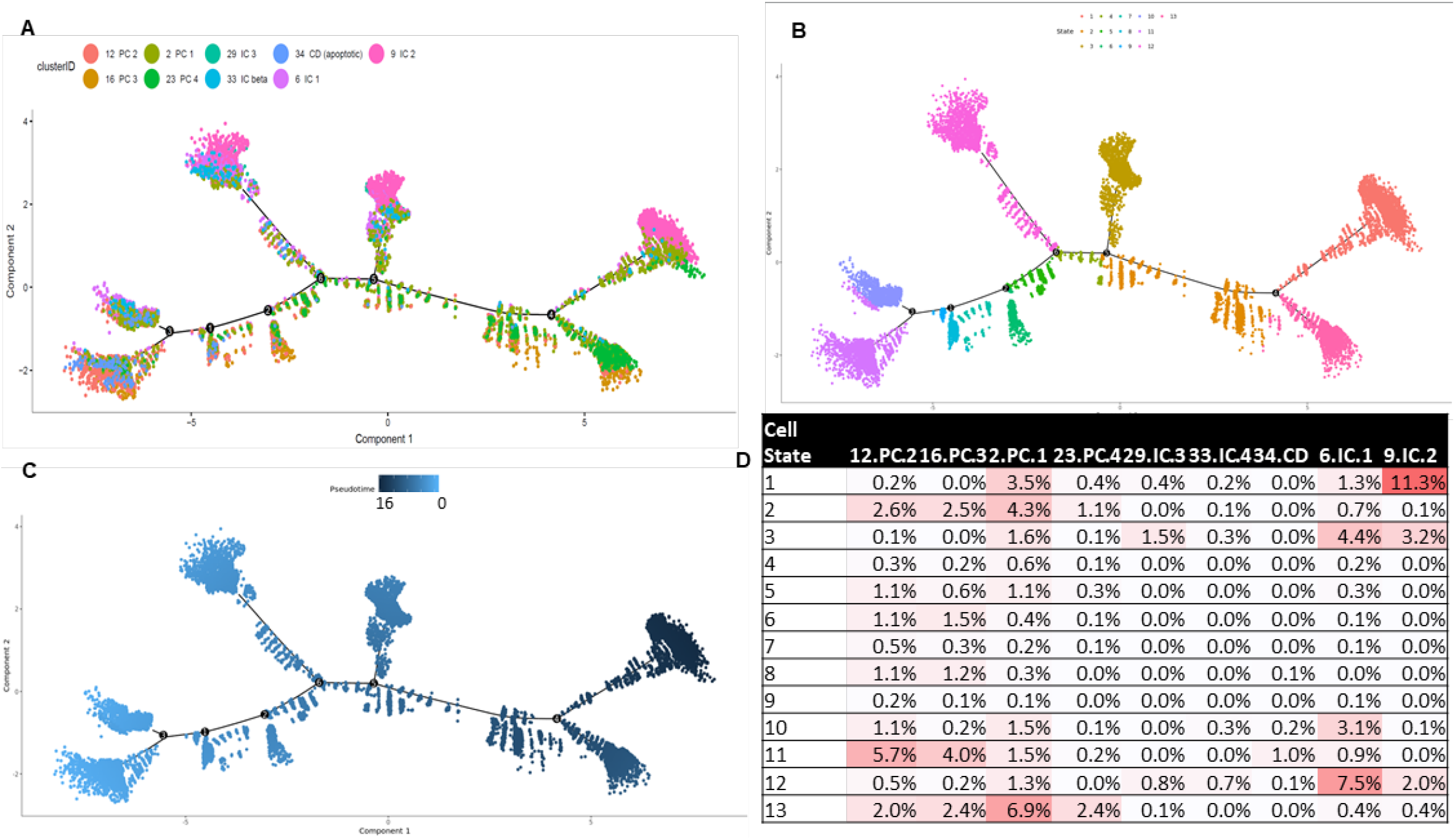
Pseudotime trajectory plots of collecting duct cells. Points are colored by origin of cluster ID (A), pseudotime-derived cell state (B), and pseudotime on a continuous scale (C). (D) Represents the proportion of all 11,326 CD cells from each cluster in (A) assigned to each cell state in (B).

**Figure S4.**
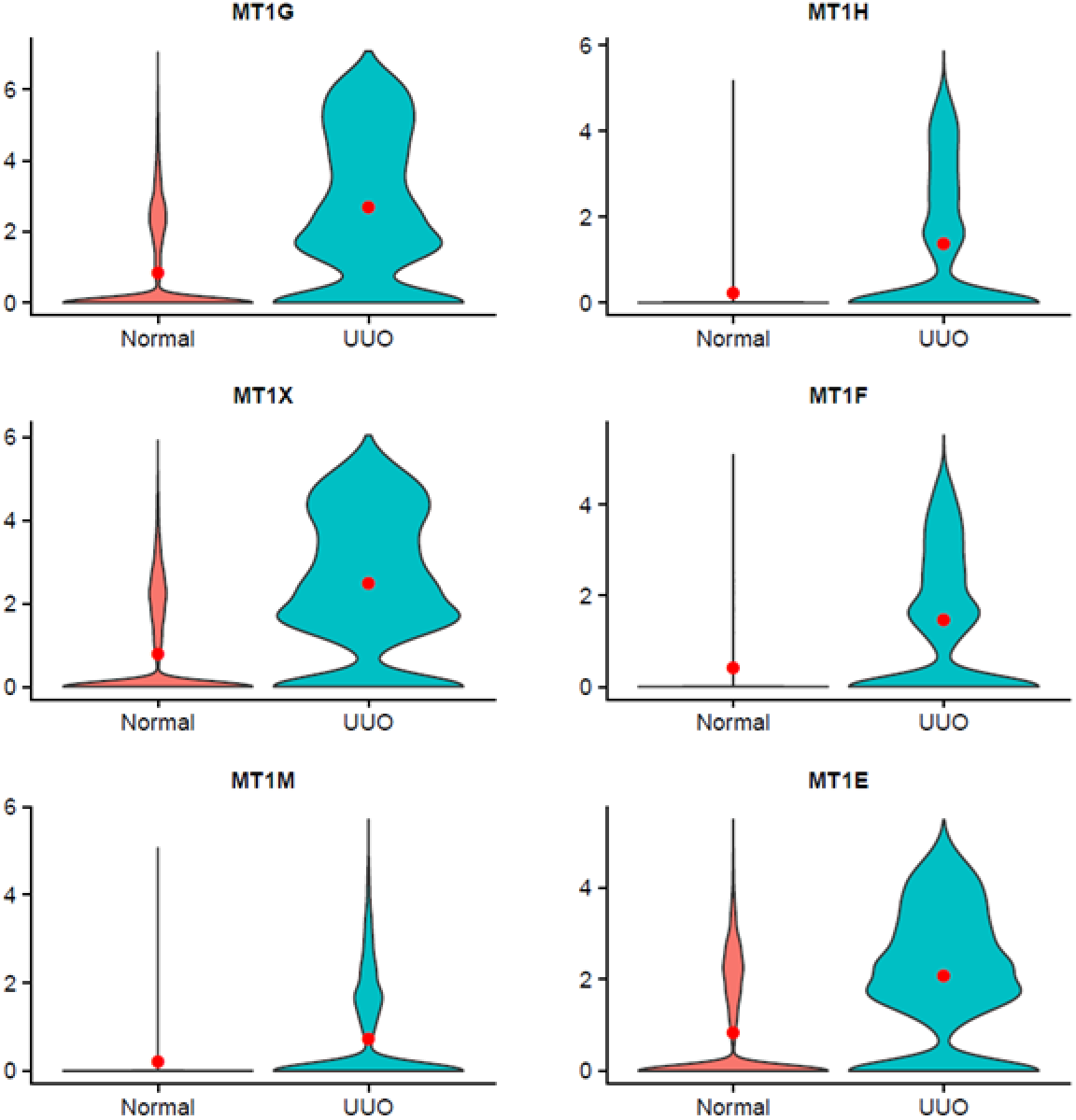
Upregulation of metallothioneins in Unilateral Ureteral Obstruction (UUO) nephrectomy. Violin plots showing distribution of metallothionein protein subunit encoding genes and their differential expression between kidney cells from tumor nephrectomies (N=40,173) and kidney cells from a UUO nephrectomy (N=5,238). Red dot overlaid on the plot signifies the mean expression value. All comparisons are equally statistically significant by Wilcoxon test at p<0.0001.

**Figure S5.**
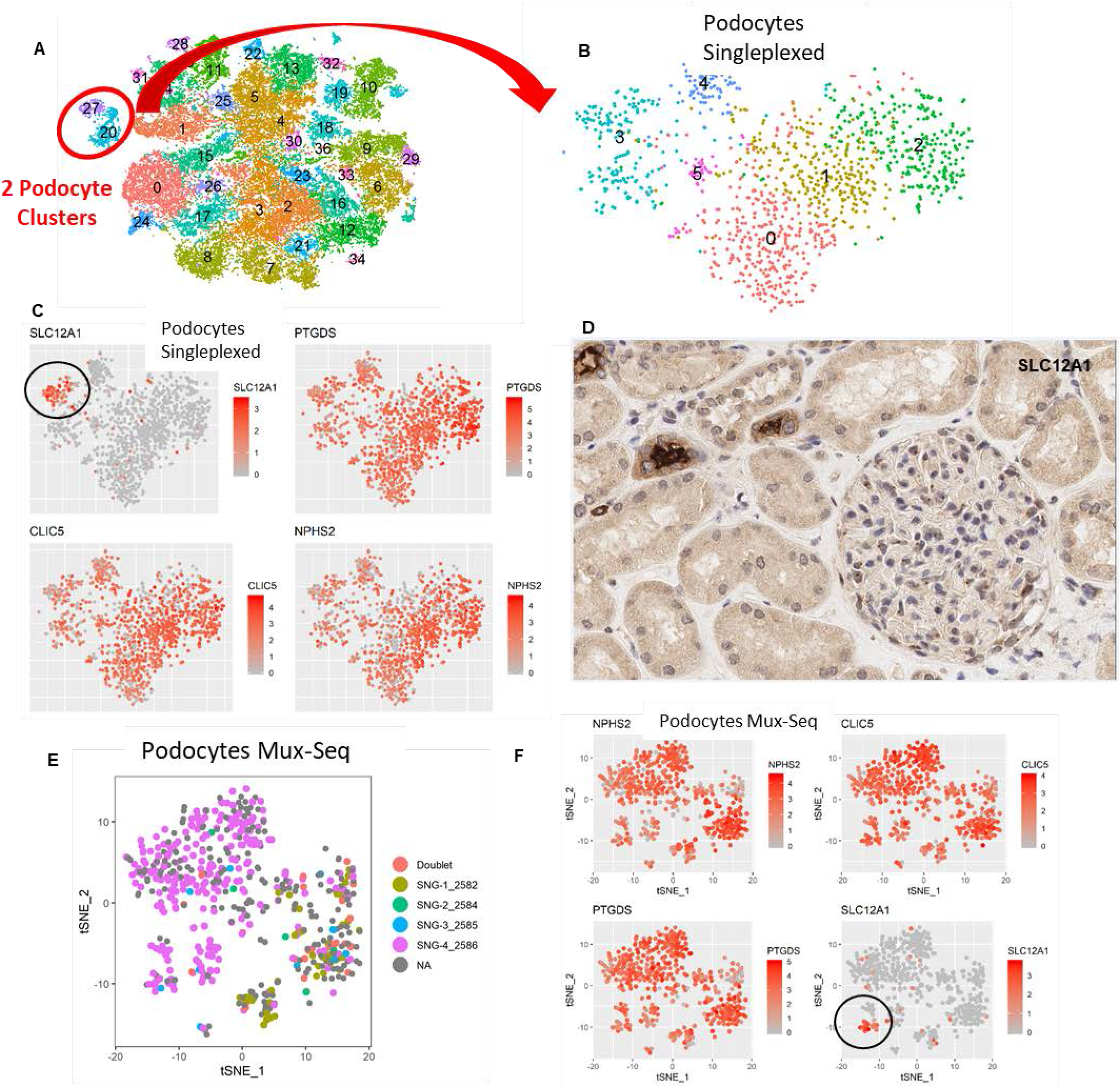
Sub cluster analysis of podocytes. (A) tSNE plot showing two podocyte clusters used for downstream sub clustering. (B) tSNE plot showing 6 distinct podocyte sub clusters. (C) tSNSE plots show three known podocyte markers expressed throughout the sub clusters, with previously known TAL marker *SLC12A1* expressed in a subset of podocytes. (D) IHC staining in renal tissue from a tumor nephrectomy shows presence of the NKCC2 protein encoded by *SLC12A1* in the glomerulus. These results are also displayed by the Mux-Seq method, with a demultiplexed tSNE in (E) showing which Podocytes were mapped to sample and (F) showing selected podocyte markers along with *SLC12A1*. The podocytes in (F) expressing SLC12A1 were not found to be doublets via Mux-Seq in (E).

**Figure S6:**
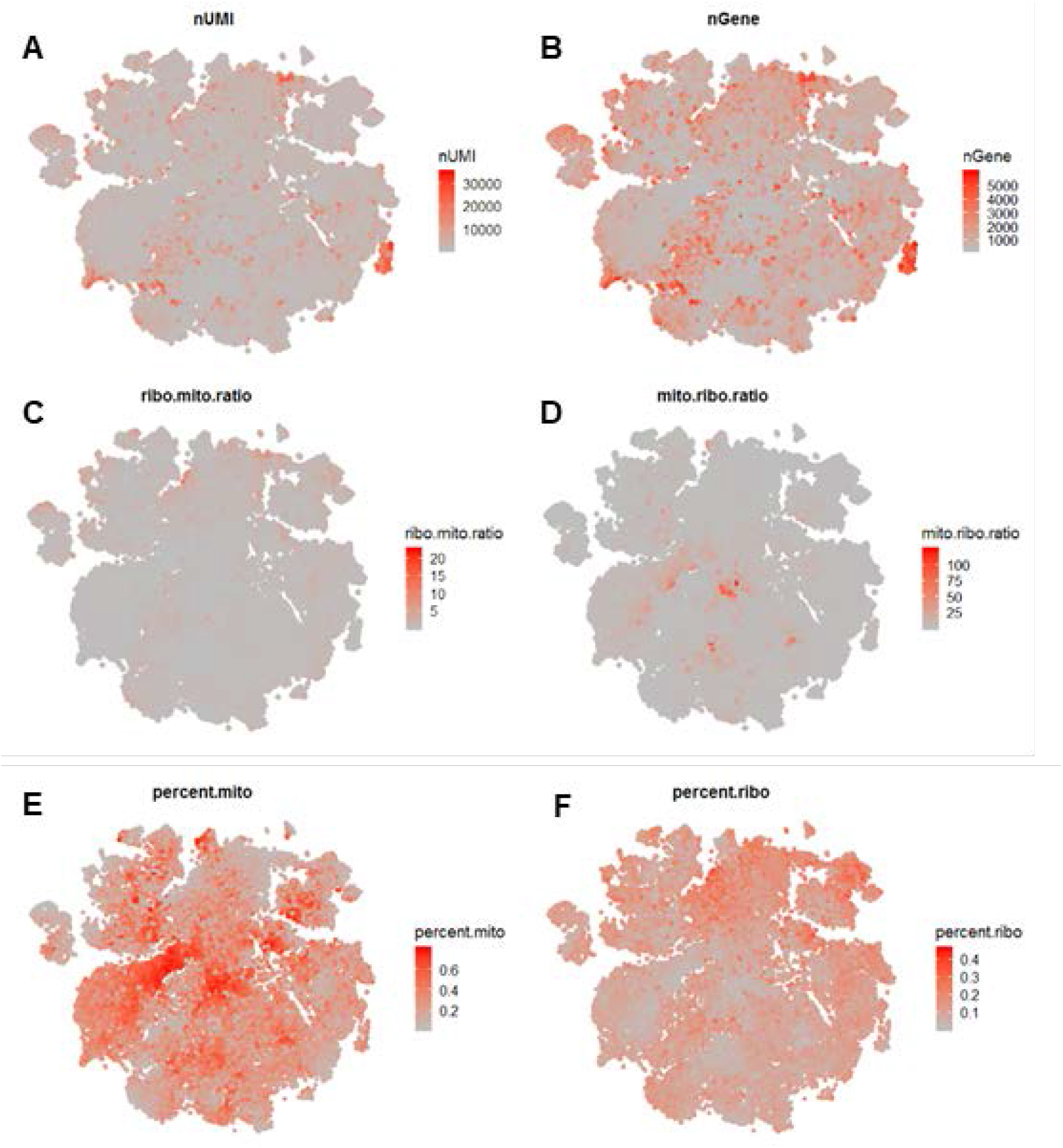
Quality control (QC) statistics of cells. All tSNE plots (A-F) represent the same cells from figure 2A but represented by six different QC metrics. These metrics are number of unique molecular identifiers per cell (nUMI) (A), number of genes per cell (nGene) (B), ratio of ribosomal to mitochondrial content (ribo.mito.ratio) to show highly proliferative cells (C), ratio of mitochondrial to ribosomal content (mito.ribo.ratio) to show stressed or apoptotic cells (D), percent mitochondrial content of whole transcriptome (percent.mito) (E), and percent ribosomal content of whole transcriptome (percent.ribo) (F)

